# Inhibition of ADAM17 increases cytotoxic effect of cisplatin in cervical spheroids and organoids

**DOI:** 10.1101/2024.01.27.577547

**Authors:** David Holthaus, Christoph Rogmans, Ina Gursinski, Alvaro Quevedo-Olmos, Marzieh Ehsani, Mandy Mangler, Inken Flörkemeier, Jörg P. Weimer, Thomas F. Meyer, Nicolai Maass, Dirk O. Bauerschlag, Nina Hedemann

**Author notes:** Shared first authorship (David Holthaus, Christoph Rogmans). Corresponding author Dr. Nina Hedemann.

## Abstract

**Background:** Cervical cancer represents one of the main causes of female, cancer-related mortality worldwide. The majority of cancers are caused by human papillomaviruses such as HPV16 and HPV18. As chemotherapeutic resistance to first-line platinum treatment is still a predominant clinical challenge in advanced cervical cancer, novel treatment options including combinatorial therapies are urgently required to overcome chemotherapeutic resistance. Inhibition of *A Disintegrin And Metalloproteinase (ADAM)*-family members, heavily involved in tumour progression of a vast range of solid tumours, strongly improved response to chemotherapeutic treatment in other tumour entities including ovarian cancer.

**Methods:** We established two- and three-dimensional models derived from three traditional cervical cancer cell lines and ectocervical cancer-derived organoids. Following characterisation, these models were used to investigate their response to cisplatin treatment in the absence and presence of ADAM inhibitors using viability assays and automated live cell imaging.

**Results:** The pivotal role of the metalloprotease ADAM17 driving chemotherapy resistance was detectable in all ectocervical cultures irrespective of the model system used, whereas ADAM10 inhibition was predominantly effective only in loosely aggregated spheroids. We showed prominent differences regarding treatment responses between 2D monolayers compared to 3D spheroid and 3D organoid model systems. Particularly, the organoid system, regarded as the closest representation of primary tumours, exhibited reliably the combinatorial effect of ADAM17 inhibition and cisplatin in all three individual donors.

**Conclusions:** As two- and three-dimensional models of the same cell lines differ in their responses to chemotherapy it is essential to validate treatment strategies in more advanced model systems representing the patient situation more realistically. Ectocervical organoids showed reliable results regarding treatment responses closely mimicking the primary tumours and could therefore serve as an important tool for personalized medicine in cervical cancer. These findings strengthen the role of ADAM17 as a potential novel target for combinatorial treatments to overcome chemoresistance in cervical cancer.

## Introduction

According to the latest WHO data, one in four female gynaecological cancer deaths worldwide can be attributed to cervical cancers [1]. Cervical carcinoma is highly associated with persistent infections with oncogenic, high-risk human papilloma viruses (HR-HPV) [2], especially HPV16 and HPV18 [1, 2]. While surgical removal of the early-stage tumour is the standard of therapy, advanced stages require radio- and chemotherapy. Since only around 25% of patients respond adequately to existing chemotherapies [3], new strategies for the treatment of cervical cancer cases are urgently needed.

Platinum resistance mechanisms in cervical cancer are manifold and include amongst others the reduced uptake and enhanced efflux of chemotherapeutics, increased DNA repair, and deactivation of pathways leading to apoptosis [4]. In other cancers, it has been shown that the A Disintegrin and Metalloproteinase (ADAM) gene family is implicated in chemotherapeutic resistance and the inhibition of family members is beneficial to improve platinum therapy *in vitro* [5–8]. The ADAM gene family is involved in a variety of biological processes by activation of several proteins such as growth factors via cleavage of membrane-bound precursor proteins [5, 6, 9]. In cancer, ADAM10 and ADAM17 have been the most actively studied [10]. Both are structurally and functionally related to each other. ADAM10s role in cancer progression and initiation is less well understood but is thought to be related to increased cell migration and invasion [10]. Elevated levels of ADAM10 have been associated with poor survival in cervical cancer [11]. In contrast, ADAM17 has already been identified as a major mediator of therapy resistance and prognosis in cancer [5, 6, 8, 12, 13]. In pathophysiological conditions, ADAM17 enhances the cleavage of growth factor ligands such as amphiregulin (AREG) and heparin-binding EGF-like growth factor (HB-EGF) [12]. We have reported an increase in ADAM17 activity and substrate release in response to cisplatin treatment in ovarian cancer cell lines and spheroids [5, 6]. Others reported that the expression of ADAM17 is associated with aggressive progression and poor prognosis in cervical cancer [14].

The translatability of traditional two-dimensional models of cancer have been debated recently [15], as those models lack features inherent of 3D tissues and organoids such as polarization and compartmentalization. A common three-dimensional alternative model is the spheroid model, composed of cancer cell lines or primary cells that aggregate in suspension or in extracellular matrices [5, 15]. Spheroid models are often used to start off with, as they are rather simple to implement and provide robust readouts [5]. Nevertheless, the arrangement of cells is more or less random and phenotypes of spheroids are predominantly cell line specific. Thus, some cell lines grow as dense spheroids including necrotic core formation and generate nutrition and penetration gradients, whereas others form rather irregular loose cell aggregates lacking this zonal compartmentalisation. In order to form more structured and differentiated structures closely representing the original tissue or tumours, a strong focus was laid on generation of more complex models such as organoids. Recent developments have enabled the culture of healthy cervical and cervical cancer organoids [16–18]. These organoids represent an *in vitro* tool that preserves cellular heterogeneity and recapitulates tissue architecture and functionality [16, 17]. It has been shown for various entities that the culture of resected tumour material as organoids accurately represents the nature of the tumours [19–21]. Initial studies show that the histological architecture of the cancer tissue, genetic signature, tumour heterogeneity and thus also therapy response can be well mapped [19, 22]. Furthermore, we have earlier established three cervical cancer organoid lines [17, 18] and shown that these lines elicit a differential response to γδ T cells in co-culture compared to healthy cervical organoids [16]. Few other studies using organoids have been presented for cervical cancer [16, 19, 23] and an *in vitro* comparison of mono- and combination treatments, for example with ADAM inhibitors, are currently missing.

To ensure an intra cell line-specific comparison whether chemotherapeutic responses are different in diverse cervical cancer models. We established two- and three-dimensional models derived from three traditional cervical cancer cell lines and ectocervical cancer-derived organoids. Additionally, we developed a robust and reliable image-based readout system for quantification of matrix-embedded individual organoids, representing a powerful tool to assess therapeutic efficacy. Following characterisation, these models were used to investigate their individual response to cisplatin treatment in the absence and presence of ADAM inhibitors using a multiplexed combination of viability assays and the above-described automated live cell analysis tool.

## Materials & Methods

### Isolation of cervical tissue and cervical organoid culture

Specimens of human ectocervix were obtained from volunteers undergoing surgery at the Department of Gynaecology, Charité University Hospital, and August-Viktoria Klinikum, Berlin (Ethics Approval EA1/059/15). Samples were processed within 3 hours after resection and informed consent was obtained from all donors. Organoids were derived from tissue resections as previously described [16–18, 24]. Organoids were passaged every 5-7 days at a 1:3-1:5 ratio and seeded in Matrigel seven days before experiments. Patient characteristics are given in Supplementary Table 1.

### Cell lines and 3T3-J2 irradiation

SIHA (ATCC, HTB-35; RRID:CVCL _0032) and CaSki (ATCC, CRL-1550; RRID:CVCL_1100) cell lines were maintained in RPMI 1640 medium supplemented with L-Glutamine, 25 mM HEPES (Capricorn, RPMI-HA), 1 mM sodium pyruvate (Capricorn, NPY-B), 1x Penicillin/Streptomycin (Gibco, 15070063) and 10% foetal calf serum (Sigma, F7524). 3T3-J2 cells (kindly provided by Craig Meyers; Howard Green laboratory, Harvard University; RRID:CVCL_W667) and C33A (ATCC, HTB-31; RRID:CVCL_1094) cell lines were maintained in DMEM supplemented with sodium pyruvate, stable Glutamine (Capricorn, DMEM-HPSTA), 1x Penicillin/Streptomycin, 10 mM HEPES and 10% foetal calf serum.

For generation of feeder cells, the 3T3-J2 cells were irradiated with 30 Gy in a Gammacell 40 Exactor. After irradiation, 1x10^6^ irradiated 3T3-J2 were seeded per T25 Flask and incubated overnight until all cells attached to the surface.

### Whole mount immunofluorescence assays (IFAs) and microscopic analyses

Phase contrast and brightfield images were taken using an IX50 (Olympus) microscope. Images were contrast adjusted and scale bars were added using FIJI [25]. For fluorescent microscopy, organoids were processed as described before [16, 26]. Images were taken using a confocal laser scanning microscope 880 microscope (Zeiss), equipped with Plan-Apochromat 20x/0.8 M27 and analysed with ZEN blue software (v3.5) and FIJI. Antibodies and dilutions are listed in Supplementary Table 2.

### Isolation of nucleic acids and real-time quantitative-polymerase chain reaction (RT-qPCR)

HPV status was assessed as described before [16]. Genomic DNA (gDNA) was isolated with the Quick-DNA Miniprep (Zymo, D3024) according to manufacturer’s instructions.

RNA was extracted from cells using Direct-zol-RNA-Microprep kit (Zymogen, R2063) including on-column DNase-I treatment following manufacturer’s protocol. RNA (>300 ng) was reverse transcribed using the Lunascript RT Supermix Kit (NEB, E3010).

RT-qPCR was performed with a StepOnePlus (Agilent) using the Luna Universal qPCR Master Mix (NEB, M3003), and included initial enzyme activation for 3 minutes at 95 °C, followed by 40 cycles of 20 s at 95 °C, 30 s at 60 °C and 20 s at 72 °C. A minimum of 5 ng cDNA/20 ng gDNA was used per well. Melting curve analysis was performed to verify amplicon specificity. Relative expression was calculated using the ΔCT method. HPV status was assessed by electrophoresis of the amplified products. Amplicons were separated on a 1.5% agarose gel containing SYBR Safe Nucleic Acid Gel Stain in 0.5x Tris-borate-EDTA-buffer. Signal was recorded using a ChemiDoc MP Imaging System (Biorad).

**Table 1:**
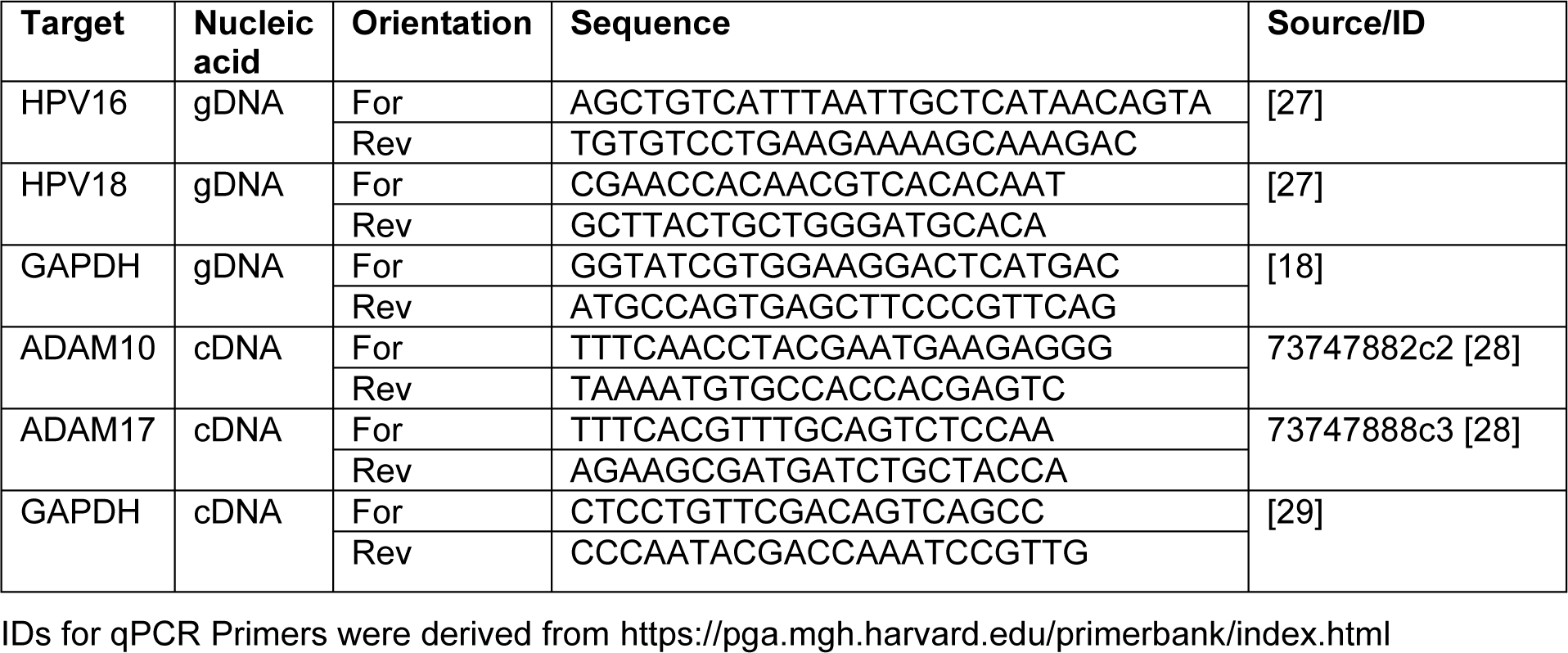
Primer sequences.

### Chemotherapeutic treatment and live cell imaging of spheroids and organoids

The imaging protocols were adapted from previous studies [5, 16]. For Spheroids, 5,000 SIHA, 7,000 CaSki and 10,000 C33A cells were seeded in ultra-low attachment (ULA), black- transparent 96-well plates (Corning, 4520). An overview of tested cell numbers is given in Supplementary Figure 1. After 24 (CaSki) to 96 (C33A and SIHA) hours of growth, cells were incubated with 0.5x CellTox Green cytotoxicity dye (Promega, G8741), cisplatin (obtained from the Clinical Pharmacy Services, UKSH, Campus Kiel), 3 µM ADAM10 inhibitor GI 254023X (Aobious, 3611) and/or 3 µM ADAM10/17 inhibitor GW 280264X (Aobious, 3632) [30, 31] or the same volume of DMSO solvent control. A positive control of 2 µg/ml of Puromycin (InVivoGen, ant-pr-1) was included. Live Cell Imaging measurements were performed using the CELLAVISTA 4 automated cell imager in combination with the SYBOT X-1000 with CYTOMAT 2 C-LiN system (all SYNENTEC). Wells were imaged every six hours for a total of 48 hours and afterwards fluorescence data and images were analysed and subsequently extracted with the YT-Software (SYNENTEC) using the Spheroid Count (2F) application. The following settings were used: Exciter: Blue (475/28) - Emissionfilter: Green Filter 530nm (530/43). In short, the application detects the spheroids in the brightfield image, and analyses the average fluorescence intensity (background-corrected) within this spheroid mask.

Cervical cancer organoid lines were pre-cultured for seven days in Collagen I pre-coated T25 flasks. Cells were then enzymatically digested with 1 ml TrypLE, pelleted at 300 xg, in Advanced DMEM/F-12 and resuspended in 1 ml of Matrigel at a concentration of 50.000 cells/ml. Cells were seeded in 10 µl Matrigeldomes in Costar Flat White Clear Bottom 96-well plates (Corning, 3903). The Matrigel was allowed to polymerize for 30 min at 37 °C and 100 µl of pre-warmed cervical organoid medium was added to the wells. The ∼500 single cells per Matrigeldome resulted in the formation of 50-100 organoids. After growth of 7 days, the organoids were treated and imaged similarly to the spheroids by addition of 0.5 x CellTox Green cytotoxicity dye with cisplatin, 3 µM ADAM10 inhibitor GI254023X and/or 3 µM ADAM10/17 inhibitor GW280264X or DMSO in 100 µl of pre-warmed cervical organoid medium. Wells were imaged every six hours for a total of 96 hours and afterwards fluorescence data and images were analysed and subsequently extracted with the YT-Software using the Spheroid Quantification (2F) application. The settings were modified to detect all organoids in the bright field channel and subsequently, the average intensity of the green channel (background-corrected) within each single organoid was analysed.

### Caspase and viability assays

For two-dimensional cell lines, a multiplexed caspase/viability assay was performed as described before [5] using the Multiplex Assay ApoLive-Glo (Promega, G6411) kit.

For spheroids and organoids, the CellTiter-Glo 3D Cell Viability Assay was performed following manufacturer’s instructions. After live cell imaging, excessive supernatant from the plates was removed. To quantify viability, the reagent was thawed overnight and allowed to equilibrate for 30 min at room temperature (RT). After equilibration, the reagent was mixed 1:1 with PBS and pipetted onto the wells. Well contents were mixed and allowed to stabilize at RT for 30 min. Afterwards, luminescence was detected using an Infinite M200 Pro or Sparks plate reader (Tecan).

### SDS-PAGE and Immunoblotting

For SDS-PAGE analysis, cells were harvested, pelleted, and washed with ice-cold PBS. For organoids, Matrigel was removed by incubation with cell-recovery solution (Corning) for 1 hour at 4 °C. All cells were lysed in RIPA-buffer (50 mM Tris-HCl pH 8.0, 1 % NP-40 (Sigma, 492018), 0.5 % sodium deoxycholate (Sigma, D6750), 0.1 % SDS (Carl Roth, 0183), 150 mM NaCl, and 5 mM EDTA). After assessing the protein content using the Pierce Rapid Gold BCA Protein Assay Kit (Thermo, A53225) samples were diluted with 6 x Laemmli buffer (60% Tris-HCl pH 6.8, 10 % SDS, 30 % glycerol (Carl Roth, 3783) and 0.01 % Bromophenol blue (Sigma, B0126) containing 10 % β-mercaptoethanol (Carl Roth, 4227). Samples were boiled at 95 °C for 5 minutes. 10 µg of protein per condition was blotted onto Amersham Protran nitrocellulose membranes (Fisher Scientific, 10600016) using standard techniques and transfer quality was validated by Ponceau S (Sigma, P3504) total protein staining. The membranes were blocked for 1 hour in 1 x RotiBlock (Roth, A151) and incubated with primary antibodies overnight. After three washing steps, membranes were incubated with respective peroxidase-conjugated secondary antibodies and signals were detected using chemiluminescence with a ChemiDoc MP Imaging System (Biorad). Antibodies and dilutions are listed in Supplementary Table 2.

### Statistical analysis and panel composition

Basic calculations were performed using MS EXCEL 2019 (Microsoft). Figures were plotted using Prism 10.1.2 (GraphPad) or R 4.1 P values ≤ 0.05 were considered as statistically significant. Asterisks indicate statistical significance values as follows: * p < 0.05, ** p < 0.01, *** p < 0.001, **** p < 0.0001. The panel composition and annotations were created using Affinity Designer 2.1.1 (Serif) or R 4.1.

## Results

### ADAM17 inhibition sensitizes cervical cancer cell lines to cisplatin treatment

For initial two-dimensional experiments, we selected three commonly used cervical cancer cell lines; C33A, CaSki, and SIHA cells. To confirm HPV integration in all lines, they were tested for HPV16 and HPV18 integration, the two most common HR-HPVs. C33A cells were confirmed negative for both viruses, while CaSki and SIHA cells were both positive for HPV16 (Figure 1A) as described before [32]. Afterwards, all cell lines were subjected to a titration with cisplatin in the presence and absence of the ADAM10 inhibitor GI254023X (GI) or the ADAM10/17 inhibitor GW280264X (GW). After two days of incubation, viability and cell death quantified by Caspase 3/7 activity were assessed (Figure 1B & C). All three cell lines reacted differentially to the treatment. SIHA cells were strongly resistant to cisplatin with an IC50 of 17.37 µM, C33A showed intermediate resistance with an IC50 of 9.595 µM, while CaSki cells were rather sensitive to cisplatin indicated by an IC50 of 3.943 µM (Figure 1B). When treating the cells with a combination of ADAM10 inhibitor GI in combination with cisplatin, IC50s increased in all cell lines (12.81 µM, C33A; 6.679 µM, CaSki; 23.84 µM, SIHA), indicating that inhibition of ADAM10 in 2D monolayers rather increased their resistance potential to cisplatin. In contrast, inhibition of both proteases using the combinatorial ADAM10/17 inhibitor GW strongly enhanced cellular response to cisplatin exhibited by significantly, decreased IC50 values (1.388 µM, C33A; 0.173 µM, CaSki; 10.52 µM, SIHA)(Figure 1B). In addition, Caspase 3/7 activity was significantly increased by more than 2-fold in all cell lines after ADAM10/17 inhibition in comparison to cisplatin monotreatment in concentrations > 15 µM (Figure 1C), whereas sole inhibition of ADAM10 by GI did not affect the response to cisplatin. Taken together, the three cell lines showed cell line-dependent responses to cisplatin treatment. Combination treatment with GW sensitized cells to cisplatin treatment, while same treatment with GI increased cell viability.

**Figure 1:**
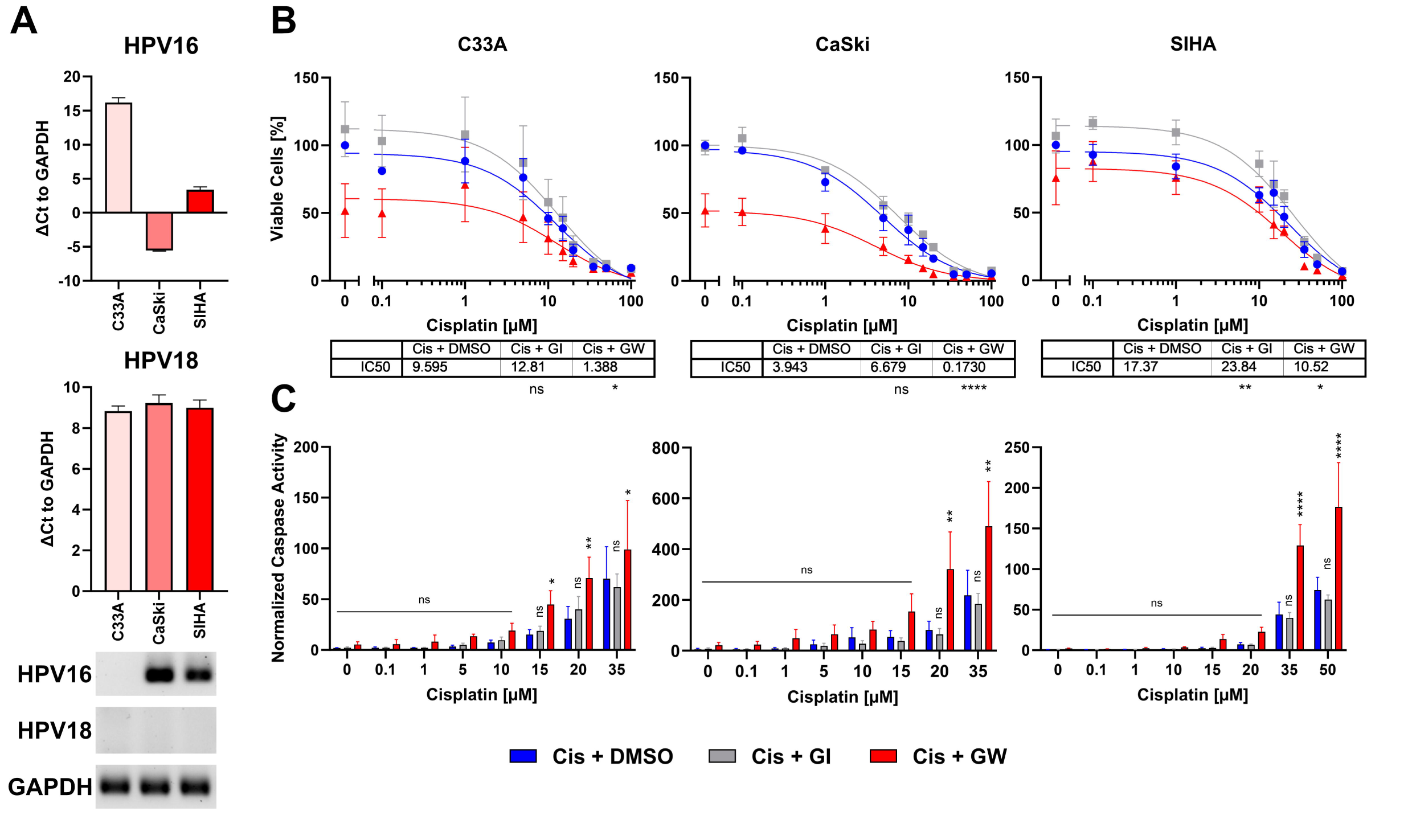
Combinatorial Effect of ADAM10/17 inhibition on two-dimensional monolayers A) Characterization of HPV integration status of C33A, CaSki, and SIHA cells. RT-qPCR experiments show mean (± SEM) from 3 independent experiments. **B)** Viability and **C)** Caspase activity quantification of C33A, CaSki, and SIHA cells after treatment with Cisplatin with/without 3 µM ADAM10 inhibitor GI254023X or ADAM10/17 GW280264X inhibitor for 48 hours. Data shows mean (± SEM) of ≥ 3 independent experiments per cell line. Statistical significance to DMSO solvent control was determined using a Two-Way-ANOVA with Tukey’s correction for multiple testing. ns not significant * p < 0.05, ** p < 0.01, *** p < 0.001, **** p < 0.0001.

### Characterization of cervical spheroids

As the translatability of two-dimensional models to *in vivo* responses is debated [15, 33], we implemented advanced three-dimensional cell culture models to validate our treatment effects (Figure 2). Cell lines grown as 3D cultures in ultra-low attachment plates showed different spheroid morphologies; while C33A and SIHA cells aggregated loosely, CaSki cells formed compact spheroids rapidly as described previously (Figure 2A, B, Supplementary Figure 2) [34]. To assess potential changes of 2D vs 3D cultures we assessed protein expression of multiple markers for either tissue origin or cancer characteristics, like Marker of Proliferation KI67 (KI67), Cytokeratin 5 (KRT5), Tumour Protein 53 (P53), and ADAM17 (Figure 2C). Interestingly, all spheroid (3D) cultures showed increased expression of KI67 in contrast to traditional 2D culture. Only CaSki cells expressed Cytokeratin 5 (KRT5), indicating ectocervical origin. Interestingly, despite being also derived from squamous epithelium, C33A and SIHA cells showed no KRT5 expression, as was pointed out before [35, 36]. P53 expression was high in C33A spheroid culture and low in both types of CaSki cultures. No significant p53 expression was detected in SIHA cells as described before [32]. We detected expression of ADAM17 in all cervical cancer cell lines and abundance of the pro-form (P) of ADAM17 was comparable between lines and culture conditions, while the active form (A) was expressed more prominently in the spheroid conditions. To assess the level of transcriptional expression between 2D and 3D we performed RT-qPCR of ADAM10 and ADAM17 (Figure 2D, Supplementary Figure 2). The overall number of transcripts was comparable. However, CaSki cells expressed significantly more transcripts after spheroids generation, while no significant difference was found in C33A and SIHA cells (Figure 2D, Supplementary Figure 2).

**Figure 2:**
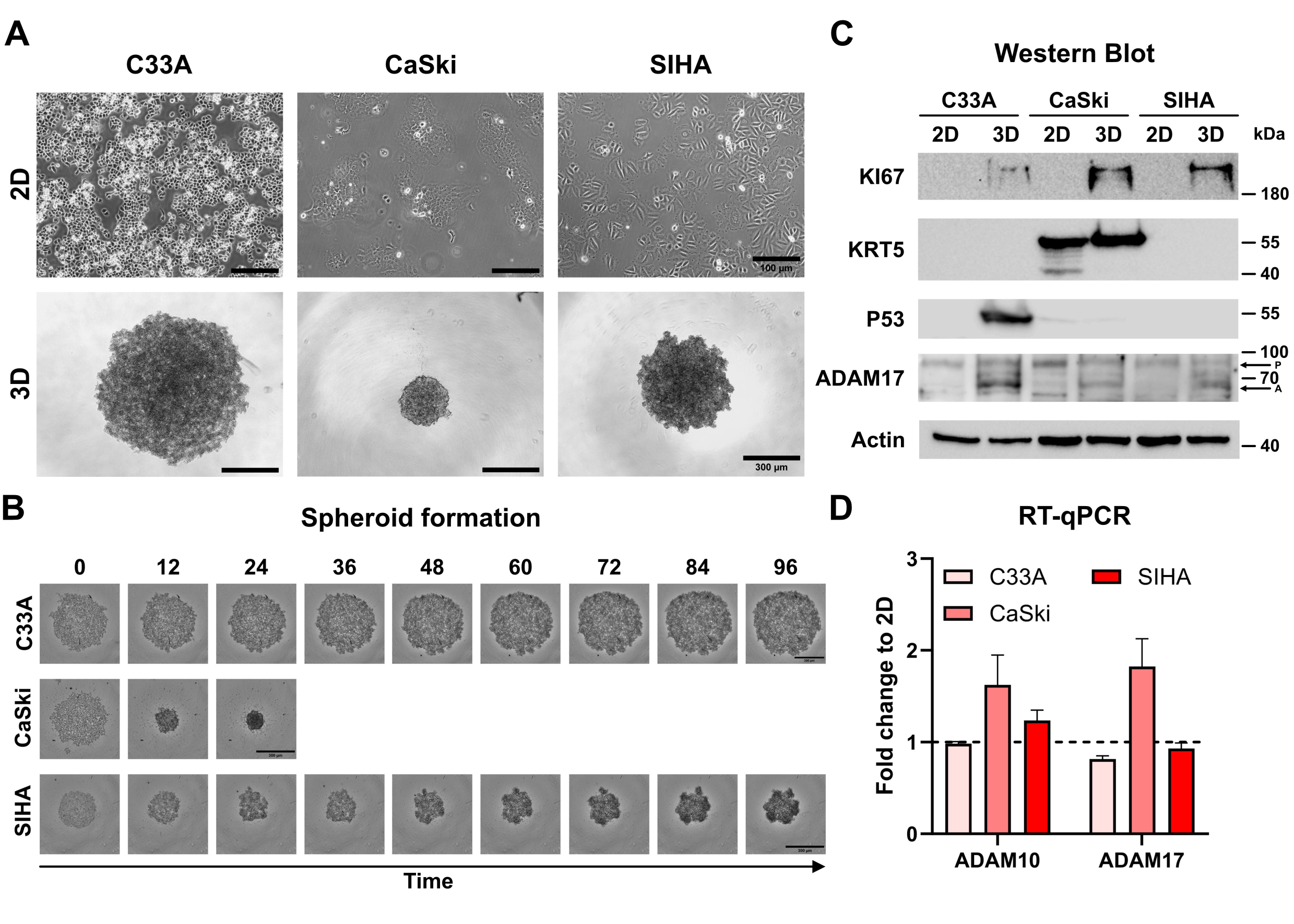
Characterization of C33A, CaSki, and SIHA spheroid cultures. A) Brightfield images of two- and three-dimensional cultures of cervical cell lines. C33A and SIHA cells show formation of loose aggregates in contrast to CaSki cells that form dense spheroids. **B)** Time course of spheroid aggregation pattern of individual cell lines. Aggregation of CaSki spheroids was completed after 24 hours. **C)** Comparison of protein expression of marker genes between two- and three-dimensional cultures by western blot. P indicates ADAM17 pro-form, and A indicates the active form. **D)** Comparison of transcriptional expression of ADAM10 and ADAM17 by RT-qPCR. Experiments show mean (± SEM) of 3 independent experiments. Scale bars indicate 100 µm (2A, upper panel) or 300 µm (2A, lower panel and 2B).

### C33A and CaSki spheroids are sensitized to cisplatin by ADAM10 and ADAM17 inhibition

To assess whether the transformation into spheroids changes the response to treatment, we subjected the spheroid cultures of all three cell lines to cisplatin treatment in the presence and absence of ADAM inhibitors. Viability was assessed by ATP quantification and live cell imaging. During the 48 hours long incubation, the spheroids were imaged every 6 hours with an automated imaging system in the presence of CellTox Green dye to stain dead cells. The live cell imaging revealed that the cell lines reacted cell line-dependent to cisplatin treatment (Figure 3). ADAM17 inhibitor GW significantly increased cytotoxicity in CaSki cells, while in C33A and SIHA cells no differences between conditions were observed (Figure 3A, Supplementary Figure 3). Quantification of cytotoxicity for all three cell lines and cisplatin concentrations is shown in Supplementary Figure 4.

**Figure 3:**
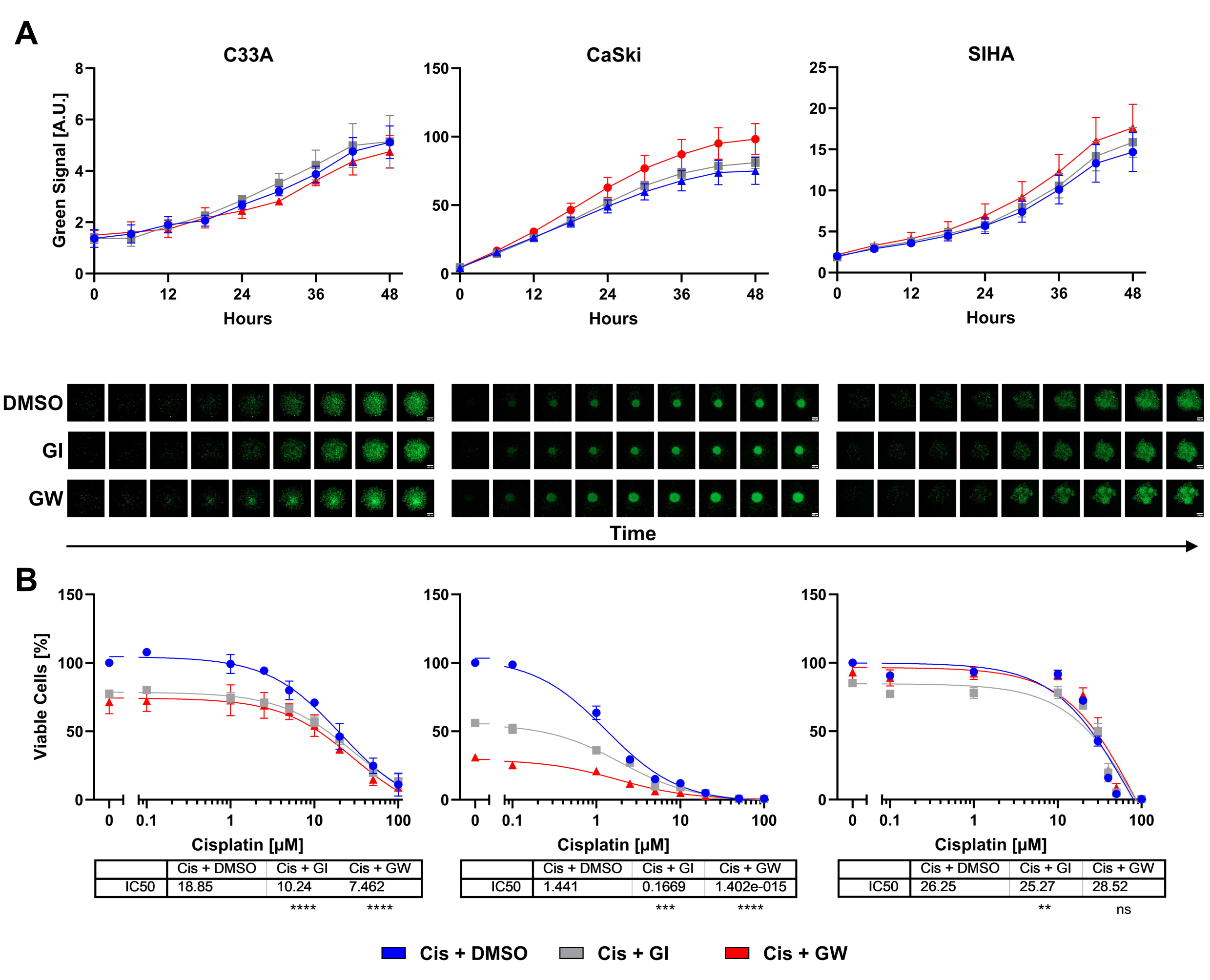
Combinatorial effect of ADAM inhibition on three-dimensional cervix cancer cell spheroid cultures. A) Accumulation of cytotoxicity as measured by live cell imaging with CellTox Green over time. Exemplary cisplatin concentrations with/without 3 µM ADAM10 inhibitor GI254023X or ADAM10/17 GW280264X inhibitor are shown; 10 µM for C33A, 1 µM for CaSki and 20 µM for SIHA cells. Data shows mean (± SEM) from 3 independent experiments per cell line. Representative images are displayed below. Scalebar indicates 100 µm. Enlarged images are given in Supplementary Figure 3. Other cisplatin concentrations are given in Supplementary Figure 4. **B)** Viability quantification of C33A, CaSki, and SIHA spheroids after treatment with Cisplatin with/without 3 µM ADAM10 inhibitor GI254023X or ADAM10/17 GW280264X inhibitor for 48 hours. Data shows mean (± SEM) of 3 independent experiments per cell line. Statistical significance to DMSO solvent control was determined using a Two-Way ANOVA with Tukey’s correction for multiple testing. ns not significant ** p < 0.01, *** p < 0.001, **** p < 0.0001.

After live cell imaging, cell viability was assessed by quantification of ATP. Responses of spheroids were similar to two-dimensional cultures (Figure 3B). CaSki cells remained highly sensitive to cisplatin. C33A and SIHA cells exhibited comparably high IC50s of 18.85 µM and 26.25 µM cisplatin, respectively (Figure 3B). In contrast to 2D cultures, GI sensitized cells to cisplatin treatment. Inhibition of ADAM10 nearly halved the IC50 for C33A and almost by a factor of 10 for CaSki cells. The combined inhibition of ADAM10 and ADAM17 was even more effective in both spheroid cultures. Interestingly, while C33A and CaSki cells showed a strong response to ADAM10 and ADAM17 inhibition, even in the absence of cisplatin, SIHA cells were not affected by ADAM inhibition as changes in both conditions were negligible (Figure 3B).

### Characterization of cervical cancer organoids as patient-specific model

To better depict tissue architecture and tumour heterogeneity in comparison to cancer cell lines, we have established organoids from cervical cancer before [16–18]. All lines could be propagated in collagen-coated cell culture flasks or extracellular matrices, like Matrigel, and were positive for ectocervical marker KRT5 (Figure 4A). Two of the lines, Pat1 and Pat2 show integrated HPV16, while Pat3 is positive for HPV18 (Figure 4B). We again assessed differences in protein expression between 2D and 3D cultures. Interestingly, all patient lines showed differences between 2D and 3D culture (Figure 4C). KI67 and P53 were only detectable in 3D cultures in all lines. Pro-forms of ADAM17 were comparable between cultures. The active form of ADAM17 showed differential expression in both 2D to 3D culture, and between patient samples, with 3D culture generally showing less expression (Figure 4C). KRT5 displayed multiple bands in 2D culture in Pat1 and Pat3, and in both culture methods of Pat2, indicating patient specific differences. To confirm expression differences of ADAM10 and ADAM17 we also conducted transcriptomic analysis by RT-qPCR (Figure 4D, Supplementary Figure 2). We could confirm the decrease of ADAM10 and ADAM17 expression at the transcriptomic level. ADAM10 and ADAM17 were significantly decreased in all patient cells after converting from 2D to 3D, except for ADAM10 in Pat3 (Figure 4D, Supplementary Figure 2). However, ADAM10 expression was also lowest in Pat3 (Supplementary Figure 2).

**Figure 4:**
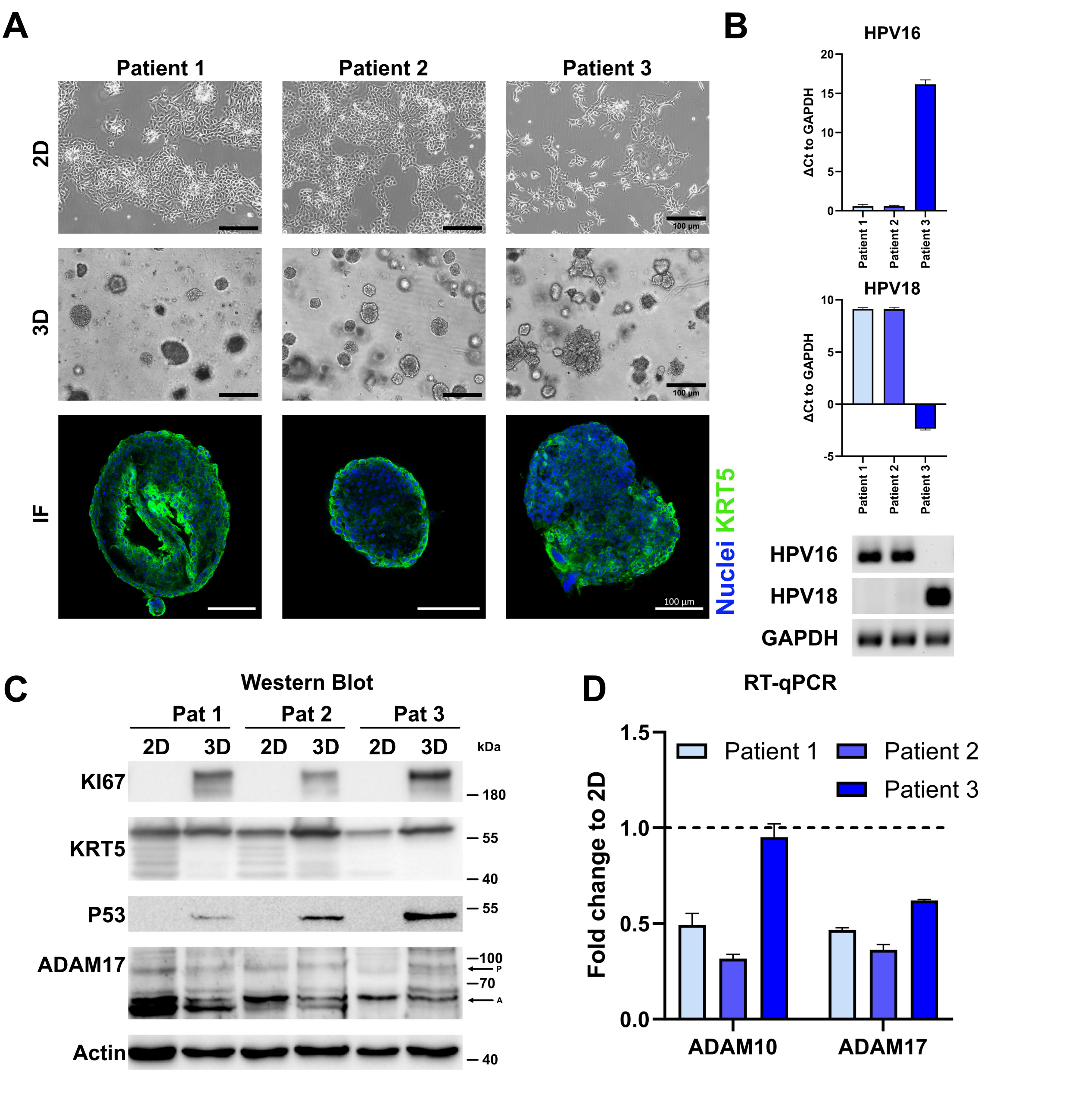
Characterization of cervical cancer organoids. A) Representative fluorescent images of three-dimensional and brightfield images of two- and three-dimensional cultures of ectocervical cancer-derived organoids. Ectocervical marker cytokeratin 5 (KRT5) is stained in green, nuclei are stained with Hoechst33342 (blue). Scale bars indicate 100 µm. **B)** Characterization of HPV integration status of all three organoid lines. RT-qPCR experiments show mean (± SEM) from 3 independent experiments. **C)** Comparison of protein expression of marker genes between two- and three-dimensional organoid cultures by western blot. P indicates ADAM17 pro-form, and A indicates the active form. **D)** Comparison of transcriptional expression of ADAM10 and ADAM17 by RT-qPCR. Data shows mean (± SEM) of 3 independent experiments.

### Cervical cancer organoids are sensitized to cisplatin by ADAM17 inhibition

To investigate whether these lines can model patient-specific differences to chemotherapeutics, we modelled the responses to cisplatin monotherapy and combinatorial treatment with ADAM inhibitors by live cell imaging (Figure 5). In comparison to untreated controls, cisplatin treated organoids showed increased cytotoxicity (Figure 5A. Supplementary Figure 3 and 4). Combinatorial inhibition of ADAM10 or ADAM17 with cisplatin significantly increased cytotoxicity in all patient lines. Following live cell imaging, again viability assays were performed. In these, all our organoid lines showed IC50s ≥ 14 (Figure 5B). Pat2 showed the highest IC50 with 26.21 µM (Figure 5B). Combinatorial treatment with the ADAM10 inhibitor GI sensitized all three different organoid lines to cisplatin. While in organoids from Pat1 the IC50 was decreased from 15.36 µM to 10.86 µM, it was reduced from 26.21 µM to 18.23 µM in Pat2, and nearly halved for Pat3 from 14.61 µM to 7.809 µM (Figure 5B). Importantly, when incubated with the ADAM10/17 inhibitor GW the IC50s further decreased. Pat1’s IC50 for cisplatin halved from 15.36 to 6.33 during combinational treatment (Figure 5B). Pat2 which showed high resistance to cisplatin monotreatment displayed a reduction of the IC50 from 26.21 to 8.443 µM cisplatin (Figure 5B), resulting in a comparable level to Pat1. Pat3-derived organoids exhibited high sensitivity to ADAM17 inhibition as monotreatment with GW already reduced viable cells by 40%. The IC50 in combinational treatment was reduced by around 90% from 14.61 µM to 1.323 µM (Figure 5B).

**Figure 5:**
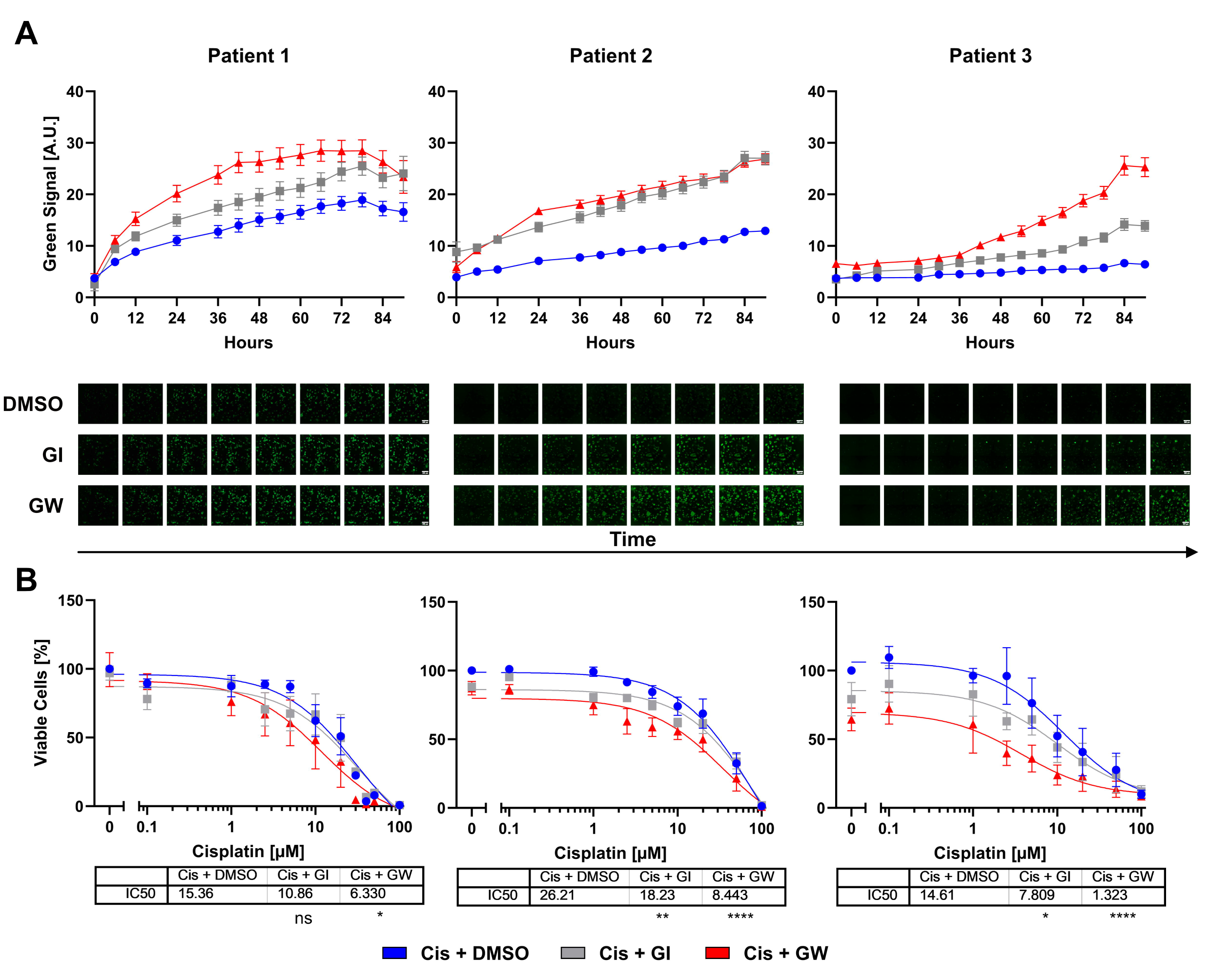
Cervical organoids show patient-specific responses to monotherapy with cisplatin and combinational treatment with ADAM inhibitors. A) Accumulation of cytotoxicity measured by live cell imaging with CellTox Green over time. A representative cisplatin concentration (10 µM) with/without 3µM ADAM10 inhibitor GI254023X or ADAM10/17 inhibitor GW280264X is displayed. Data shows mean (± SEM) from individual organoids of ≥ 3 independent experiments per patient. Up to 243 organoids per condition were assessed. Exemplary images are displayed below. Scalebar indicates 300 µm. Enlarged images are given in Supplementary Figure 3. Other cisplatin concentrations are given in Supplementary Figure 4. B) Viability quantification of single organoids after treatment with Cisplatin with/without 3 µM ADAM10 inhibitor GI254023X or ADAM10/17 GW280264X inhibitor for 96 hours. Data shows mean (± SEM) of ≥ 3 independent experiments per patient. Statistical significance to DMSO solvent control was determined using a Two-Way ANOVA with Tukey’s correction for multiple testing. ns not significant * p < 0.05, ** p < 0.01, **** p < 0.0001.

In summary, combinational treatment using ADAM10/17 inhibitors strengthened the effect of cisplatin in ectocervical organoids. Cervical cell lines differed in their responses to chemotherapy between two- and three-dimensional models of the same cell lines. Inhibition of ADAM17 showed a potent effect on cisplatin sensitivity in all models and culture conditions. These findings strengthen the role of ADAM17 as a potential novel target for combinatorial treatments to overcome chemoresistance in cervical cancer.

## Discussion

During the past decades, the importance of advanced patient-derived model systems, particularly for cancer therapy, is continuously growing. Here, we present a comparison of multiple cellular model systems to study novel combinatorial treatments to overcome chemotherapy resistance in cervical cancers focusing on the inhibition of the metalloproteases ADAM10 and ADAM17. Even though we showed prominent differences regarding treatment responses between two-dimensional monolayers compared to three-dimensional spheroid- and three-dimensional organoid model systems, the pivotal role of the metalloprotease ADAM17 driving chemotherapy resistance was detectable in all cultures irrespective of the model system used. Particularly, the organoid system, regarded as the closest representation of primary tumours [21], presented reliably the combinatorial effect of ADAM17 inhibition and cisplatin in all three individual patients. These findings strengthen the role of ADAM17 as a potential novel target for combinatorial treatments to overcome chemoresistance in cervical cancer.

Alike other cancers, chemoresistance in cervical cancer presents one of the major clinical challenges [4]. Therefore, the generation of novel treatments or treatment combinations together with or besides chemotherapy and radiation is fundamentally needed. Recently, antiangiogenic agents like bevacizumab and the checkpoint inhibitor pembrolizumab (Clinical trial NCT04221945)[3] were added to the standard platinum-paclitaxel-based first line chemotherapy, leading to an increase in progression-free survival in advanced cervical cancer. Other strategies include the combination of platin-based therapeutics with PARP inhibitors, HPV vaccinations, and Ribonucleotide reductase (RNR) inhibitors [37]. Nevertheless, the rate of non-responders is still substantial [38].

The metalloproteases ADAM10 and ADAM17 have been studied extensively in the past decades [39]. Next to the regulation of inflammatory processes they play an essential role in tissue regeneration and pathologically in tumour development. Malignant cellular behaviour, such as invasion, proliferation, and angiogenesis, was associated with enhanced expression or activation of these ADAM-members *in vitro* and resulted in enhanced tumour growth *in vivo* [10]. Tissue expression of ADAM17 correlated with severe outcomes in mammary carcinoma patients and underlined its critical role in cancer [40]. Most of the pathological effects of these proteases in relation to cancer can be explained by an enhanced growth factor release like Amphiregulin (AREG), heparin binding-EGF like growth factor (HB-EGF) or epithelial growth factor (EGF). By binding to their respective receptors like epithelial growth factor receptor (EGFR), Erb-B2 Receptor Tyrosine Kinase 2 (ERBB2/HER2), HER3 and other receptor tyrosine kinases. Activation of well-known downstream cascades like the MAP Kinases or Phosphatidylinositol-4,5-Bisphosphate 3-Kinase (PI3K)/ AKT Serine/Threonine Kinase (Akt) finally leads to enhanced tumour cell survival, inhibition of apoptotic signalling or enhanced metastasis [10, 39].

In cervical cancer, infection with HR-HPV plays an essential role in disease initiation and consequently, almost all cervical cancers are HPV-positive, mostly with HPV16 and HPV18 [2]. The oncogenicity is mainly instigated by subunits E6 and E7 that cause Tumour protein 53 (TP53) degradation, inhibition of Cyclin Dependent Kinase Inhibitor 1A (CDKN1A/P21) and RB Transcriptional Corepressor 1 (RB1), and E2F Transcription Factor (E2F) activation leading to continuous cell cycle activation and decreased apoptosis [41]. In head and neck cancer, it has been described that HPV-negative and -positive cancers differ in deviant pathway activation and their responses to chemotherapeutics [42]. While in cervical cancer HPV-positive cancer represents the majority, responses to chemotherapy differ to HPV-negative cancers mainly due to differences in mutational signatures [43]. In contrast to head and neck cancer, is it believed that HPV-positive cervical cancer is less chemoresistant as DNA damage repair is altered due to genetic HPV-induced changes in pathways involving, amongst others, TP53, Cyclin Dependent Kinase Inhibitor 2A (CDKN2A/p16), and Lysine Acetyltransferase 5 (KAT5/TIP60) [44]. In contrast to chemotherapy, we have earlier demonstrated that HPV- oncogene E6E7-transformed healthy cervical cells, and HPV-positive cervical cancer cells exhibit higher cytotoxicity mediated by γδ T cells when in co-culture [16]. Interestingly, also ADAM10 and 17 have been shown to be induced by HPV infection [11, 45]. Next to the well- known pro-tumorigenic characteristics of ADAM17, it was demonstrated, that in cervical cancer, expression of ADAM17 is actually required for the HPV entry platform assembly via CD9, transforming growth factor alpha (TGFα) and phosphorylated Mitogen-Activated Protein Kinase 1 (MAPK1/ERK) [45]. Therefore, and based on our prior study focussing chemoresistance in ovarian cancer we aimed to unravel a potential mechanism of chemotherapy associated activation of these proteases in cervical cancer.

The role of ADAMs in chemotherapy resistance has been described earlier for other entities such as cervical cancer, liver cancer, colorectal cancer and bladder cancer [5–8, 10, 46]. In line with our previous studies focussing on chemoresistance in ovarian cancer [5–7], we showed strong combinatorial effects of ADAM17 inhibition and cisplatin treatment in cervical cancer. Interestingly, these effects were most prominent in two-dimensional cultures, three-dimensional ectocervical cultures of CaSki cells as well as in primary ectocervical organoids, but less pronounced in spheroids of SIHA and C33A. Differences in treatment responses in 2D vs. 3D have been reported frequently and can be explained by various mechanisms leading from differential target expression patterns to differences in diffusion rates through media, outer cell layers or extracellular matrices [47]. For our particular set of cell lines, others showed similar divergences in treatment responses [48]. In line, with our data on SIHA and C33A, it has been reported that tumour spheroids show elevated chemoresistance in comparison to two-dimensional cultures [49]. Based on current literature, tight spheroids generally tend to represent rather chemoresistant phenotypes compared to loosely formed aggregates. We observed this phenomenon in our OvCa spheroids likewise, but not in our cervical model [5]. Although CaSki cells, formed tight spheroids, they responded with higher cytotoxicity to cisplatin treatment compared to loose aggregate forming SIHA and C33A cells [49]. Therefore, we hypothesise that in our model setup there are other more dominant factors affecting chemoresistance apart from hypoxic gradients and diffusion of chemotherapeutics, which underscores the effects of 3D organization in the biology of cells.

Surprisingly, from the three cell lines, only CaSki cells expressed KRT5, a marker for ectocervix. However, all of them are described to be derived from squamous cell carcinoma. While endo- and ectocervical cancers differ in their mutational signatures and behaviour, the absence of essential markers of tissue origin indicates significant deviations from initial tissue-derived programming that could influence resistance mechanisms and cellular behaviour. These changes are indicative of the differences in junctional protein expression between cell lines. Supporting this, SIHA and C33A have been shown to lack expression of major cell-to-cell junctional compartments such as E-Cadherin (CDH1) [50]. Additionally, CaSki showed distinct cellular invasion patterns compared to SIHA cells that invade significantly slower [34]. It can be speculated that inhibition of ADAM10 is more effective in 3D spheroid cultures as ADAM10 plays a major role in cell migration and invasion, and thus in restructuring of cell-to-cell connections. Supporting this, cleavage of CDH1, N-Cadherin (CDH2), L1 Cell Adhesion Molecule (L1CAM), and CD44 by ADAM10 has been reported [10]. While both, ADAM10 and ADAM17, have been shown to modulate cell-to-cell and cell-to-matrix interactions [10, 51], differential expression was not uniform between cell lines and patients after conversion from 2D to 3D. Further studies should investigate other ADAM family members that have been implicated in the observed processes by others [52].

Until recently, cervical cancer modelling using organoids has been largely underrepresented in comparison to other entities, such as ovarian carcinomas [19, 53, 54]. While cancer organoid protocols share similarities, most include varying cocktails of growth factors and hormones, such as epithelial growth factor (EGF), WNT Family Member ligands, and/or Hepatocyte Growth Factor (HGF). It is likely that specific culture conditions favour specific mutational loads and the importance of culture conditions on chemoresistance and marker gene expression has been pointed out recently [54]. So far, few other groups have described the establishment of cervical cancer organoids [23, 55, 56]. In our study, patient-derived organoids show clear deviations from immortalized cell lines. All of our isolates show resistant phenotypes that are affected by ADAM10/17 inhibition. Interestingly, while all our organoid isolates exhibited resistant phenotypes to cisplatin treatment, others found varying responses based on tumour origin and phenotype [55, 56].

Several groups and companies focussed on the generation of ADAM17 targeting antibodies or ADAM17 inhibitors like INCB7839 (summarized in Wang, Xuan [46]), of which some reached Phase II trials. Most of those studies had to be terminated because of side effects or lack of drug efficacy. Clinical studies focussing on ADAM17 inhibition often suffer from high toxicity and high structural homology of the catalytic domains of ADAM-family members. Recently the focus has shifted on the non-catalytic domains that appear to be more promising [46, 57]. Therefore, our individualised organoid systems, complemented with our newly developed automated readout tools to predict treatment responses, would present an important tool to foster personalised therapy in cervical cancer. In hindsight of our study and the results of others [54–56], the optimization of cervical cancer organoid establishment pipelines with the aim of personalized medicine and treatment options should focussed, especially if surgery is excluded as therapeutic option. The direct correlation of driver mutations, protein expression and activation to potential therapy options should be emphasized and pave the way to novel target driven therapies. The involvement of ADAM10/17 inhibitors during treatment could enhance the chemotherapeutic efficacy of first line treatment such as cisplatin therapy.

## Declarations

### Ethics approval and consent to participate

The studies involving humans were approved by the ethical committees of the Charité Berlin (Ethics Approval EA1/059/15) and the Medical Faculty of the Christian-Albrechts-Universität zu Kiel (Ethics Approval B292/23). The studies were conducted in accordance with the local legislation and institutional requirements. The participants provided their written informed consent to participate in this study.

### Consent for publication

Not applicable

### Availability of data and materials

Data sharing is not applicable to this article as no datasets were generated or analysed during the current study.

### Competing interests

The authors declare that they have no competing interests.

### Authors’ contributions

DH, CR, IG, AQO, and ME conducted experiments. DH, CR, IG, and AQO analysed and visualized results. DH, AQO, and NH established methods and analysed experiments. MM took biopsies and provided tissues. IF and J-P.W supported analysis. TFM, NM and DOB provided infrastructure. TFM, DOB, NM and NH provided funding. NH conceptualized and supervised the project. DH, CR, and NH wrote the draft manuscript. All authors critically interpreted the data and revised the manuscript for important intellectual content. All authors contributed to the article and approved the submitted version.

## Acknowledgments

The authors thank Sigrid Hamann, Finja Grundt and Frauke Grohmann for excellent technical assistance. We thank Prof. Daniela Wesch, Institute of Immunology, UKSH Kiel for her support with 3T3-J2 feeder cell irradiation. We thank Wiebe Schaefer, Dr. Reinhild Geisen, and Dr. Anna Willms from SYNENTEC GmbH for the provision of the CELLAVISTA 4 automated cell imager, the SYBOT X-1000, the CYTOMAT 2 C-LiN system, and technical assistance throughout the live cell experiments.

## Funding

The authors acknowledge funding from the University Hospital Schleswig-Holstein. TFM acknowledges funding from BMBF Infect-ERA project “CINOCA” (FK 031A409A) and ERC Advanced grant “MADMICS” ID: 885008. ME is a recipient of the Focus Biomed Foundation.

## Notes

### Competing Interest Statement

The authors have declared no competing interest.

## References

1. World Health Organisation. Cervical cancer - Factsheet. 2023 06.12.2023]; Available from: https://www.who.int/news-room/fact-sheets/detail/cervical-cancer.

2. Okunade, K.S., Human papillomavirus and cervical cancer. J Obstet Gynaecol, 2020. 40(5): p. 602–608.

3. Lorusso, D., et al., LBA38 Pembrolizumab plus chemoradiotherapy for high-risk locally advanced cervical cancer: A randomized, double-blind, phase III ENGOT-cx11/GOG- 3047/KEYNOTE-A18 study. Annals of Oncology, 2023. 34: p. S1279–S1280.

4. Zhu, H., et al., Molecular mechanisms of cisplatin resistance in cervical cancer. Drug Des Devel Ther, 2016. 10: p. 1885–95.

5. Hedemann, N., et al., ADAM17 Inhibition Increases the Impact of Cisplatin Treatment in Ovarian Cancer Spheroids. Cancers (Basel), 2021. 13(9).

6. Hedemann, N., et al., ADAM17 inhibition enhances platinum efficiency in ovarian cancer. Oncotarget, 2018. 9(22): p. 16043–16058.

7. Hugendieck, G., et al., Chemotherapy-induced release of ADAM17 bearing EV as a potential resistance mechanism in ovarian cancer. J Extracell Vesicles, 2023. 12(7): p. e12338.

8. Kyula, J.N., et al., Chemotherapy-induced activation of ADAM-17: a novel mechanism of drug resistance in colorectal cancer. Clin Cancer Res, 2010. 16(13): p. 3378–89.

9. Dusterhoft, S., J. Lokau, and C. Garbers, The metalloprotease ADAM17 in inflammation and cancer. Pathol Res Pract, 2019. 215(6): p. 152410.

10. Mullooly, M., et al., The ADAMs family of proteases as targets for the treatment of cancer. Cancer Biol Ther, 2016. 17(8): p. 870–80.

11. Guo, X., et al., Elevated Expression of ADAM10 Induced by HPV E6 Influences the Prognosis of Cervical Cancer. Genet Test Mol Biomarkers, 2023. 27(5): p. 165–171.

12. Sahin, U., et al., Distinct roles for ADAM10 and ADAM17 in ectodomain shedding of six EGFR ligands. J Cell Biol, 2004. 164(5): p. 769–79.

13. Sharma, A., et al., Secretome Signature Identifies ADAM17 as Novel Target for Radiosensitization of Non-Small Cell Lung Cancer. Clin Cancer Res, 2016. 22(17): p. 4428–39.

14. Xu, Q., et al., ADAM17 is associated with EMMPRIN and predicts poor prognosis in patients with uterine cervical carcinoma. Tumour Biol, 2014. 35(8): p. 7575–86.

15. Kayser, C., et al., The challenge of making the right choice: patient avatars in the era of cancer immunotherapies. Front Immunol, 2023. 14: p. 1237565.

16. Dong, J., et al., γδ T cell-mediated cytotoxicity against patient-derived healthy and cancer cervical organoids. Frontiers in Immunology, 2023. 14.

17. Gurumurthy, R.K., et al., Patient-derived and mouse endo-ectocervical organoid generation, genetic manipulation and applications to model infection. Nat Protoc, 2022. 17(7): p. 1658–1690.

18. Koster, S., et al., Modelling Chlamydia and HPV co-infection in patient-derived ectocervix organoids reveals distinct cellular reprogramming. Nature Communications, 2022. 13(1): p. 1030.

19. Hoffmann, K., et al., Stable expansion of high-grade serous ovarian cancer organoids requires a low-Wnt environment. The EMBO Journal, 2020. 39(6): p. e104013.

20. Battistini, C. and U. Cavallaro, Patient-Derived In Vitro Models of Ovarian Cancer: Powerful Tools to Explore the Biology of the Disease and Develop Personalized Treatments. Cancers (Basel), 2023. 15(2).

21. Drost, J. and H. Clevers, Organoids in cancer research. Nat Rev Cancer, 2018. 18(7): p. 407–418.

22. Nanki, Y., et al., Patient-derived ovarian cancer organoids capture the genomic profiles of primary tumours applicable for drug sensitivity and resistance testing. Scientific Reports, 2020. 10(1): p. 12581.

23. Lohmussaar, K., et al., Patient-derived organoids model cervical tissue dynamics and viral oncogenesis in cervical cancer. Cell Stem Cell, 2021. 28(8): p. 1380–1396 e6.

24. Chumduri, C., et al., Opposing Wnt signals regulate cervical squamocolumnar homeostasis and emergence of metaplasia. Nat Cell Biol, 2021. 23(2): p. 184–197.

25. Schneider, C.A., W.S. Rasband, and K.W. Eliceiri, NIH Image to ImageJ: 25 years of image analysis. Nat Methods, 2012. 9(7): p. 671–5.

26. Holthaus, D., et al., Harmonization of Protocols for Multi-Species Organoid Platforms to Study the Intestinal Biology of Toxoplasma gondii and Other Protozoan Infections. Front Cell Infect Microbiol, 2020. 10(935): p. 610368.

27. Herfs, M., et al., A discrete population of squamocolumnar junction cells implicated in the pathogenesis of cervical cancer. Proc Natl Acad Sci U S A, 2012. 109(26): p. 10516–21.

28. Wang, X., et al., *PrimerBank: a PCR primer database for quantitative gene expression analysis*, *2012 update*. Nucleic Acids Res, 2012. 40(Database issue): p. D1144-9.

29. Holthaus, D., et al., Dissection of Barrier Dysfunction in Organoid-Derived Human Intestinal Epithelia Induced by Giardia duodenalis. Gastroenterology, 2022. 162(3): p. 844–858.

30. Chalaris, A., et al., Apoptosis is a natural stimulus of IL6R shedding and contributes to the proinflammatory trans-signaling function of neutrophils. Blood, 2007. 110(6): p. 1748–55.

31. Ludwig, A., et al., Metalloproteinase inhibitors for the disintegrin-like metalloproteinases ADAM10 and ADAM17 that differentially block constitutive and phorbol ester-inducible shedding of cell surface molecules. Comb Chem High Throughput Screen, 2005. 8(2): p. 161–71.

32. Srivastava, S., et al., The status of the p53 gene in human papilloma virus positive or negative cervical carcinoma cell lines. Carcinogenesis, 1992. 13(7): p. 1273–5.

33. Jubelin, C., et al., Three-dimensional in vitro culture models in oncology research. Cell & Bioscience, 2022. 12(1): p. 155.

34. Muniandy, K., et al., Growth and Invasion of 3D Spheroid Tumor of HeLa and CasKi Cervical Cancer Cells. Oncologie, 2021. 23(2): p. 279--291.

35. Uhlen, M., et al., A pathology atlas of the human cancer transcriptome. Science, 2017. 357(6352): p. eaan2507.

36. The Human Protein Atlas. https://www.proteinatlas.org. 2024 [cited 2024 18.01.].

37. Regalado Porras, G.O., . Chávez Nogueda, and A. Poitevin Chacón, Chemotherapy and molecular therapy in cervical cancer. Rep Pract Oncol Radiother, 2018. 23(6): p. 533–539.

38. Duenas-Gonzalez, A., Combinational therapies for the treatment of advanced cervical cancer. Expert Opin Pharmacother, 2023. 24(1): p. 73–81.

39. Zunke, F. and S. Rose-John, The shedding protease ADAM17: Physiology and pathophysiology. Biochim Biophys Acta Mol Cell Res, 2017. 1864(11 Pt B): p. 2059-2070.

40. McGowan, P.M., et al., ADAM-17 predicts adverse outcome in patients with breast cancer. Annals of Oncology, 2008. 19(6): p. 1075–1081.

41. Watkins, D.E., et al., Advances in Targeted Therapy for the Treatment of Cervical Cancer. Journal of Clinical Medicine, 2023. 12(18): p. 5992.

42. Mandal, R., et al., The head and neck cancer immune landscape and its immunotherapeutic implications. JCI Insight, 2016. 1(17): p. e89829.

43. Lee, J.E., et al., Untold story of human cervical cancers: HPV-negative cervical cancer. BMB Rep, 2022. 55(9): p. 429–438.

44. Spiotto, M.T., et al., Biology of the Radio- and Chemo-Responsiveness in HPV Malignancies. Semin Radiat Oncol, 2021. 31(4): p. 274–285.

45. Mikuličić, S., et al., Tetraspanin CD9 affects HPV16 infection by modulating ADAM17 activity and the ERK signalling pathway. Med Microbiol Immunol, 2020. 209(4): p. 461–471.

46. Wang, K., et al., Immunomodulatory role of metalloproteinase ADAM17 in tumor development. Front Immunol, 2022. 13: p. 1059376.

47. Kenny, P.A., et al., The morphologies of breast cancer cell lines in three-dimensional assays correlate with their profiles of gene expression. Molecular Oncology, 2007. 1(1): p. 84–96.

48. Yi, S.A., et al., HP1γ Sensitizes Cervical Cancer Cells to Cisplatin through the Suppression of UBE2L3. International Journal of Molecular Sciences, 2020. 21(17): p. 5976.

49. Han, S.J., S. Kwon, and K.S. Kim, Challenges of applying multicellular tumor spheroids in preclinical phase. Cancer Cell International, 2021. 21(1): p. 152.

50. Cunniffe, C., et al., Expression of tight and adherens junction proteins in cervical neoplasia. British Journal of Biomedical Science, 2012. 69(4): p. 147–153.

51. White, J.M., ADAMs: modulators of cell–cell and cell–matrix interactions. Current Opinion in Cell Biology, 2003. 15(5): p. 598–606.

52. Zigrino, P., et al., Adam-9 expression and regulation in human skin melanoma and melanoma cell lines. International Journal of Cancer, 2005. 116(6): p. 853–859.

53. Kopper, O., et al., An organoid platform for ovarian cancer captures intra- and interpatient heterogeneity. Nature Medicine, 2019. 25(5): p. 838–849.

54. Thorel, L., et al., Comparative analysis of response to treatments and molecular features of tumor-derived organoids versus cell lines and PDX derived from the same ovarian clear cell carcinoma. J Exp Clin Cancer Res, 2023. 42(1): p. 260.

55. Seol, H.S., et al., Preclinical investigation of patient-derived cervical cancer organoids for precision medicine. J Gynecol Oncol, 2023. 34(3): p. e35.

56. Kusakabe, M., et al., Application of organoid culture from HPV18-positive small cell carcinoma of the uterine cervix for precision medicine. Cancer Medicine, 2023. 12(7): p. 8476–8489.

57. Calligaris, M., et al., Strategies to Target ADAM17 in Disease: From its Discovery to the iRhom Revolution. Molecules, 2021. 26(4).

