## Supplementary Material for "Inhibition of ADAM17 increases cytotoxic effect of cisplatin in cervical spheroids and organoids"

* Shared first authorship

[#] **Corresponding author**

Dr. Nina Hedemann

**Supplementary methods:**

### Triple staining of spheroid cultures

To determine cell numbers for spheroid generation, 100-10,000 cells were seeded in ultra-low attachment, black-transparent 96-well plates (Corning, 4520). Cells were grown for 24 (CaSki) or 96 (SIHA and C33A) hours. Then, 5 µg/ml Hoechst33342 (Thermo Fisher, H1399), 0.2 µg/ml Propidium iodide (Miltenyi Biotec, 130-093-233), and 1 ng/ml Calcein Green (Invitrogen, C34852) was added to the cells. Following a three-hour incubation at room temperature, cells were imaged with the NyOne imaging system (SYNENTEC). Images were exported using YT software (SYNENTEC) and scale bars were added in FIJI.

### Supplementary Figures

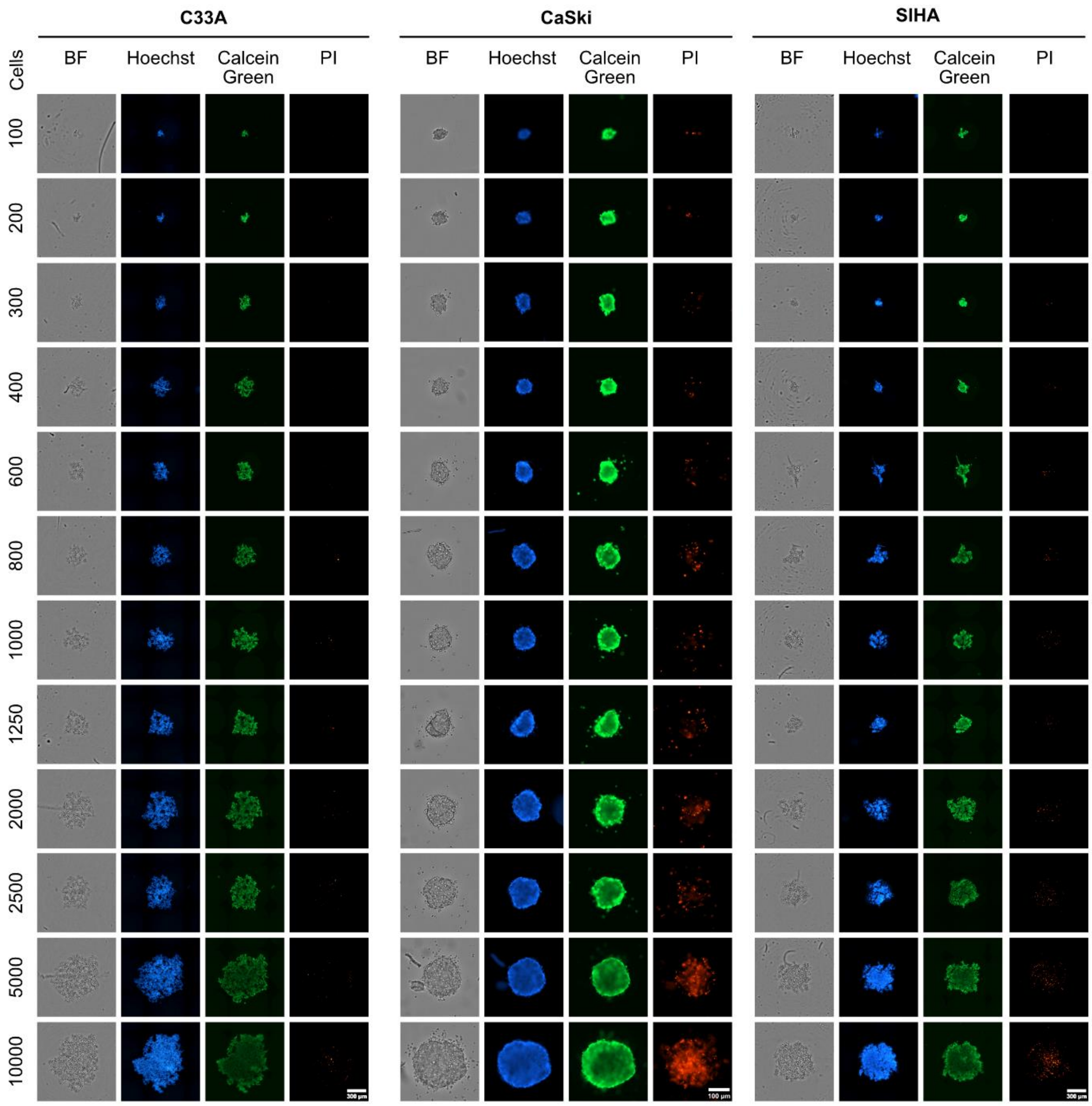


**Supplementary Figure 1: Triple staining of spheroids composed of various cell numbers of C33A, CaSki and SIHA cells.** *Cells were stained with Hoechst33342, Calcein Green and propidium iodide (PI). For later experiments, 10,000 C33A cells, 7,000 CaSki cells and 5,000 SIHA cells were used as necrotic core formation as observed by PI staining was low. Scale bars indicate 300 µm for C33A and SIHA, and 100 µm for CaSki.*

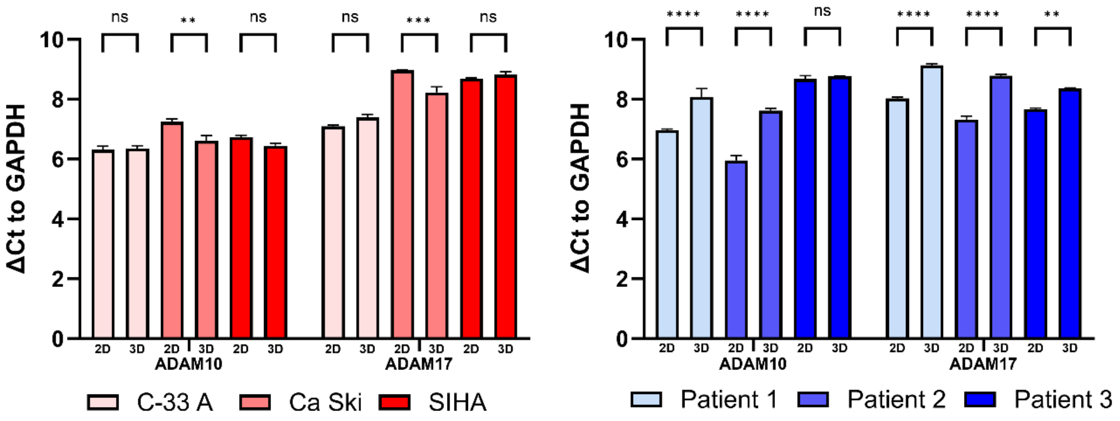


**Supplementary Figure 2: Transcriptional expression of ADAM10/17 normalized to GAPDH as measured by RT-qPCR.** *Data shows mean (± SEM) from 3 independent experiments. Statistical significance was determined using a Two-Way ANOVA with Tukey's correction for multiple testing. ns not significant * $p < 0.05$, ** $p < 0.01$, *** $p < 0.001$**** $p < 0.0001$.*

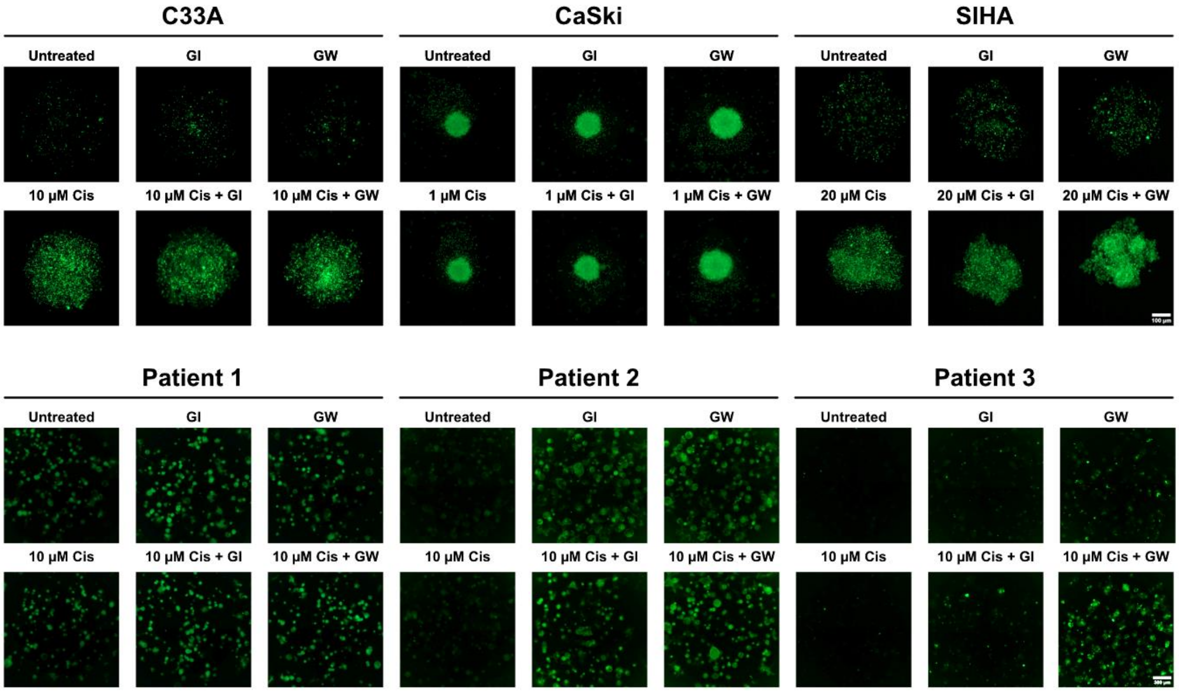


**Supplementary Figure 3: Representative images of CellTox Green staining after 36 (cell lines) or 72 (organoids) hours of incubation.** *Spheroids or organoids were of either untreated, treated with ADAM10 inhibitor GI254023X, treated with ADAM10/17 inhibitor GW280264X, or treated with combinations of cisplatin and ADAM inhibitors. CellTox green marks dead cells. Scale bars indicate 100 µm for cell lines and 300 µm for organoids.*

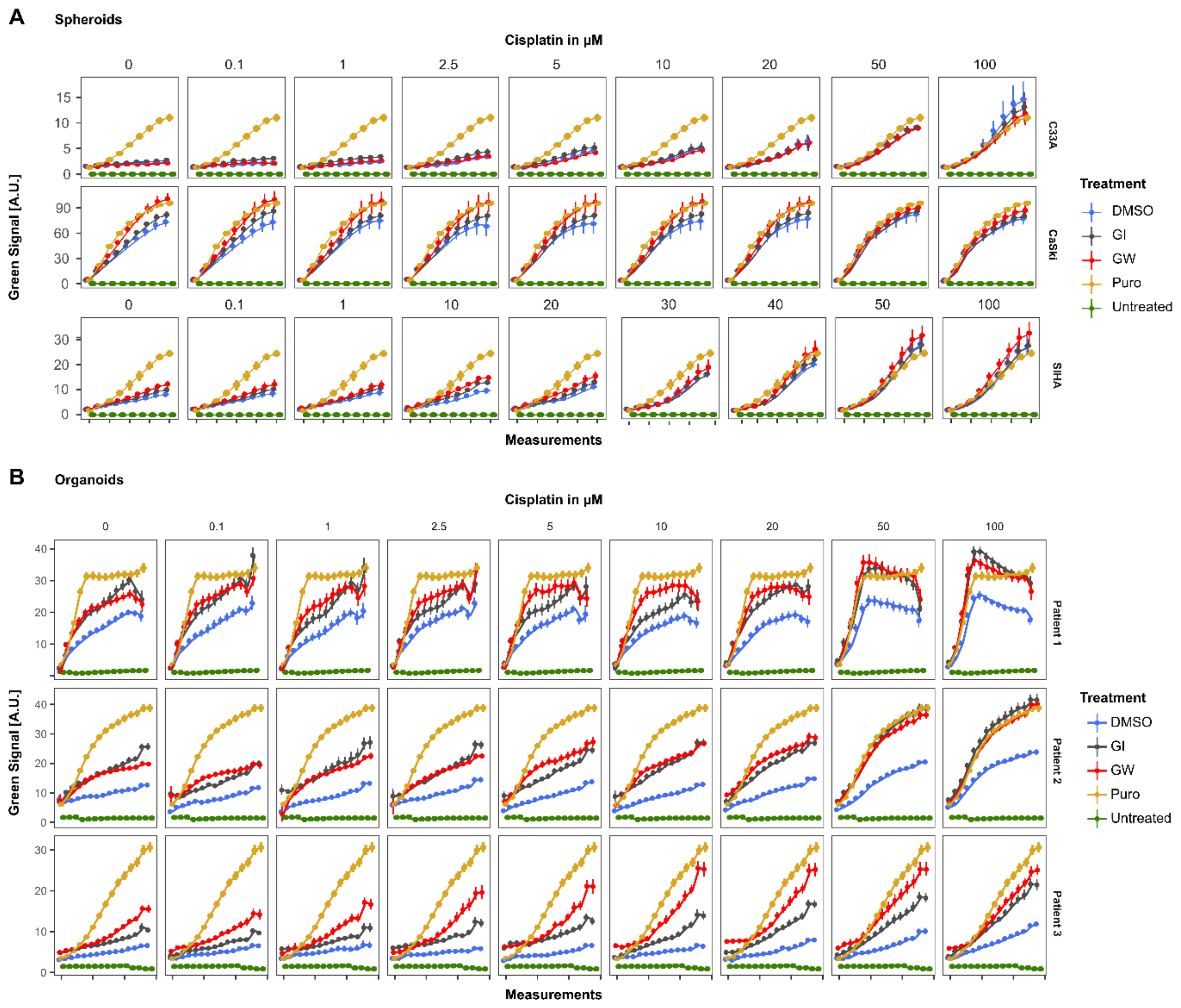


**Supplementary Figure 4: Accumulation of cytotoxicity measured by live cell imaging with CellTox Green over time.** *Varying Cisplatin concentrations with/without 3µM ADAM10 inhibitor GI254023X or ADAM10/17 inhibitor GW280264X are displayed. Data shows mean (± 95% CI) from ≥ 3 independent experiments per patient or cell line. 2 µg/ml of Puromycin was included as a positive control for cytotoxicity. Untreated controls indicate unstained and untreated organoids.*

### Supplementary tables

**Supplementary Table 1: Patient characteristics of the organoid lines.**

| ID | Age at surgery | Condition | HPV status |
|---|---|---|---|
| Patient 1 | 35 | Squamous cell carcinoma | HPV16+ |
| Patient 2 | 54 | Squamous cell carcinoma | HPV16+ |
| Patient 3 | 61 | Squamous cell carcinoma | HPV18+ |

**Supplementary Table 2: Antibodies, dyes and dilutions.**

| Target | Host | Vendor | Application | Dilution |
|---|---|---|---|---|
| **Primary antibodies** | | | | |
| KRT5-A488 | Rabbit | Abcam, ab193894, RRID:AB_2893023 | IF | 1:500 |
| KI67 | Rabbit | Abcam, ab16667, RRID:AB_302459 | WB | 1:1,000 |
| P53 | Mouse | Santa Cruz (DO-1), sc-126, RRID:AB_628082 | WB | 1:1,000 |
| ADAM17 | Goat | R&D Systems, AF9301, RRID:AB_10891879 | WB | 1:1,000 |
| ACTB | Mouse | Sigma, A5441, RRID:AB_476744 | WB | 1:10,000 |
| **Secondary antibodies** | | | | |
| Anti-Goat HRP | Donkey | Abcam, ab97110, RRID:AB_10679463 | WB | 1:3,000 |
| Anti-Rabbit HRP | Goat | Cell Signalling, 7074S, RRID:AB_2099233 | WB | 1:3,000 |
| Anti-Mouse HRP | Horse | Cell Signalling, 7076S, RRID:AB_330924 | WB | 1:3,000 |
| **Dyes** | | | | |
| Hoechst33342 | - | Thermo Fisher, H1399 | IF/Live cell imaging | 5 µg/ml |
| Propidium iodide | - | Miltenyi Biotec, 130-093-233 | Live cell imaging | 0.2 µg/ml |
| Calcein Green | - | Invitrogen, C34852 | Live cell imaging | 1 ng/ml |
| CellTox Green | - | Promega, G8741 | Live cell imaging | 0.5x, 1:2,000 |
